# Investigation of a derived adverse outcome pathway (AOP) network for endocrine-mediated perturbations

**DOI:** 10.1101/2021.09.14.460266

**Authors:** Janani Ravichandran, Bagavathy Shanmugam Karthikeyan, Areejit Samal

**Affiliations:** The Institute of Mathematical Sciences (IMSc), Chennai 600113, India; Homi Bhabha National Institute (HBNI), Mumbai 400094, India

**Keywords:** Adverse outcome pathway (AOP), AOP network, Endocrine disruption, Graph theory, Endocrine disrupting chemicals (EDCs), Risk assessment

## Abstract

An adverse outcome pathway (AOP) is a compact representation of the available mechanistic information on observed adverse effects upon environmental exposure. Sharing of events across individual AOPs has led to the emergence of AOP networks. Since AOP networks are expected to be functional units of toxicity prediction, there is current interest in their development tailored to specific research question or regulatory problem. To this end, we have developed a detailed workflow to construct a comprehensive endocrine-specific AOP (ED-AOP) network. Connectivity analysis of the ED-AOP network comprising 48 AOPs reveals 7 connected components and 12 isolated AOPs. Subsequently, we apply standard network measures to perform an in-depth analysis of the two largest connected components of the ED-AOP network. Notably, the graph-theoretic analyses led to the identification of important events including points of convergence or divergence in the ED-AOP network. Detailed analysis of the largest component in the ED-AOP network gives insights on the systems-level perturbations caused by endocrine disruption, emergent paths, and stressor-event associations. In sum, the derived ED-AOP network can be used to address the current knowledge gaps in the existing regulatory framework and aid in better risk assessment of environmental chemicals.

## 1. Introduction

The traditional toxicity testing and risk assessment methods are not commensurate with the ever-growing rate of new chemicals introduced into the commerce (Ankley et al., 2010; Ankley and Edwards, 2018; Edwards et al., 2016; Jeong and Choi, 2017). The US National Research Council published a vision report in 2007 titled ‘Toxicity testing in the 21st century: a vision and a strategy’ which contains several recommendations to improve and accelerate chemical toxicity testing (Council, 2007). To achieve rapid, efficient and cost-effective screening of chemicals, the report (Council, 2007) advocated the implementation of high-throughput screening methods such as *in vitro* toxicology (Edwards et al., 2016; Hartung, 2009; Krewski et al., 2020, 2010). Further, the report (Council, 2007) emphasized on the concept of ‘toxicity pathways’ and their use for chemical risk assessment. Essentially, ‘toxicity pathways’ were envisioned to capture the cellular response network whose perturbation upon chemical exposure leads to the observed adverse effects (Edwards et al., 2016; Hartung, 2009; Kleensang et al., 2014; Krewski et al., 2020, 2010; Vinken et al., 2017). On similar lines, Ankley *et al*. (Ankley et al., 2010) proposed a systematic framework namely, Adverse Outcome Pathways (AOPs), to capture the mechanistic information on observed adverse effects in humans or wildlife upon chemical exposure. Subsequently, several studies have reported the development of specific AOPs and their applications in risk assessment (Aguayo-Orozco et al., 2019a; Jornod et al., 2020; Knapen et al., 2018; Sakuratani et al., 2018; Sewell et al., 2018; Villeneuve et al., 2018a, 2014a, 2014b; Vinken, 2018, 2013; Vinken et al., 2017). In 2012, the Organisation for Economic Co-operation and Development (OECD) launched an international program to formalize the development and evaluation of AOPs. This has led to a series of OECD guidance documents (OECD, 2018, 2017, 2013) and primary literature (Aguayo-Orozco et al., 2019a; Jornod et al., 2020, p. 4; Knapen et al., 2018; Sakuratani et al., 2018; Sewell et al., 2018; Villeneuve et al., 2018a, 2014a, 2014b; Vinken, 2018, 2013; Vinken et al., 2017) for the development of AOPs and their potential applications in human- and eco-toxicology.

An AOP has been defined by Ankley *et al*. (Ankley et al., 2010) as: “the conceptual construct that portrays existing knowledge concerning the linkage between a direct molecular initiating event and an adverse outcome at a biological level of organization relevant to risk assessment”. Subsequently, Villeneuve *et al*. (Villeneuve et al., 2014a) have outlined five core principles for the development of AOPs which are as follows. Firstly, AOPs are not chemical-specific. Secondly, AOPs are modular and composed of reusable components, specifically, key events (KEs) and key event relationships (KERs). Thirdly, a single AOP can be viewed as a pragmatic unit of development and evaluation. Fourthly, multiple AOPs that share KEs and KERs among them can be viewed together as an ‘AOP network’, and such a network is likely to be the functional unit of prediction for most real-world scenarios. Fifthly, AOPs are live documents that can change over time with updates in existing knowledge. Further, Villeneuve *et al*. (Villeneuve et al., 2014a) have defined KEs in an AOP as: “a measurable change in biological state that is essential, but not necessarily sufficient for the progression from a defined biological perturbation toward a specific adverse outcome”. A KE in an AOP is assigned to one of different possible levels of biological organization namely, molecular, cellular, tissue, organ, individual and population (Villeneuve et al., 2014a). Among KEs, Molecular Initiating Events (MIEs) and Adverse Outcomes (AOs) are two special types of events that are anchored at the upstream end and downstream end, respectively, of an AOP (Villeneuve et al., 2014a). Specifically, MIEs capture the initial molecular level interactions between chemicals or stressors and their target receptor(s). In contrast, AOs capture perturbations at the organ or higher levels of biological organization such as changes in morphology or physiology. Lastly, a KER represents a directed relationship between a pair of KEs wherein one KE is upstream and another KE is downstream (Villeneuve et al., 2014a, 2014b; Vinken et al., 2017). Simply stated, the MIEs lead to AOs in an AOP via a series of KERs.

Since AOP development is envisaged to be a world-wide scientific community effort, OECD started the AOP knowledge base (AOP-KB; https://aopkb.oecd.org/) in 2014 to ensure international harmonization and encourage collaboration. Within AOP-KB, AOP-Wiki (https://aopwiki.org) is an actively maintained module with real-time updates that serves as a central repository of AOPs at various stages of development (Villeneuve et al., 2014a, 2014b). Besides serving as the central repository, AOP-WiKi has enabled the exploration of emergent AOP networks (wherein multiple AOPs are connected via shared KEs), for various research and regulatory applications (Knapen et al., 2018; Pollesch et al., 2019). To this end, we here derive and analyze an AOP network specific to endocrine disruption based on information in AOP-Wiki.

Network-centric approaches can aid in unraveling the organizing principles of complex biological systems (Barabási and Oltvai, 2004). A primary goal of the emerging discipline, computational systems toxicology, is to harness network and systems biology approaches in building predictive toxicological models through heterogeneous data integration across diverse levels of biological organization (Aguayo-Orozco et al., 2019b; Hartung et al., 2017, 2012; Sturla et al., 2014). The AOP framework has an inherent modular structure which enables sharing of KEs and KERs between individual AOPs, and this sharing of KEs leads to emergence of ‘AOP networks’ within AOP-Wiki as mentioned above (Knapen et al., 2018; Pollesch et al., 2019; Villeneuve et al., 2018a). Knapen *et al*. (Knapen et al., 2018) have defined an ‘AOP network’ as: “an assembly of 2 or more AOPs that share one or more KEs, including specialized KEs such as MIEs and AOs”. Recent studies (Coady et al., 2019; Hecker and LaLone, 2019; Howdeshell et al., 2017; Knapen et al., 2018, 2015; Villeneuve et al., 2018a) have illuminated the potential of such AOP networks in capturing interactions among individual AOPs, revealing undiscovered relationships among complex biological pathways, and investigating specific questions in toxicology. To date, 9 AOP networks have been derived from AOP-Wiki to address specific toxicity problems related to reproduction (Howdeshell et al., 2017; Knapen et al., 2015), development (Knapen et al., 2015), nervous system (LaLone et al., 2017; Spinu et al., 2019), liver (Arnesdotter et al., 2021), metabolism (Knapen et al., 2018; Villeneuve et al., 2018a) and immune system (Villeneuve et al., 2018b). Graph-theoretic analysis (Barabási and Oltvai, 2004) of these derived AOP networks has helped in revealing topological features, identifying critical paths, and characterizing interactions among AOPs in the constructed networks (Knapen et al., 2018; Villeneuve et al., 2018a).

Endocrine disrupting chemicals (EDCs) are an important class of toxic chemicals that mimic or block or interfere with hormonal functions of the endocrine system (Diamanti-Kandarakis et al., 2009; Gore et al., 2015; Karthikeyan et al., 2019; Zoeller et al., 2012). The risk assessment and regulation of EDCs pose several challenges including lack of standardized definition and validated testing methods (Darbre, 2015; Fuhrman et al., 2015). Endocrine disruption by environmental chemicals is a complex process that can occur via multiple and disparate mechanisms (Diamanti-Kandarakis et al., 2009; Gore et al., 2015; Schug et al., 2013; Zoeller et al., 2012). The AOP framework is ideal for organizing the existing knowledge and providing a pathway perspective on diverse modes of endocrine disruption by EDCs (Browne et al., 2020; Carvaillo et al., 2019; Rugard et al., 2020). Moreover, the development and analysis of an AOP network specific to endocrine disruption has the potential to reveal key events, critical paths, and unexpected links between individual AOPs capturing varied adverse effects (Knapen et al., 2018; Villeneuve et al., 2018a). Previously, there have been few efforts to construct AOP networks for disruption specific to a single hormone, namely, androgen (Howdeshell et al., 2017), thyroid, or thyroxine (Knapen et al., 2018; Villeneuve et al., 2018a). Due to the focus on specific hormones, the constructed AOP networks in these studies do not provide a comprehensive picture of all endocrine disruption mechanisms captured within AOP-Wiki. In this study, we therefore construct and analyze a comprehensive derived AOP network for endocrine disruption based on extensive manual curation of the information contained in AOP-Wiki (Figure 1; Methods). We expect this derived AOP network for endocrine disruption to shed light on gaps in the existing regulatory framework and aid in the development of new endpoints or assays for better risk assessment of EDCs.

**Figure 1:**
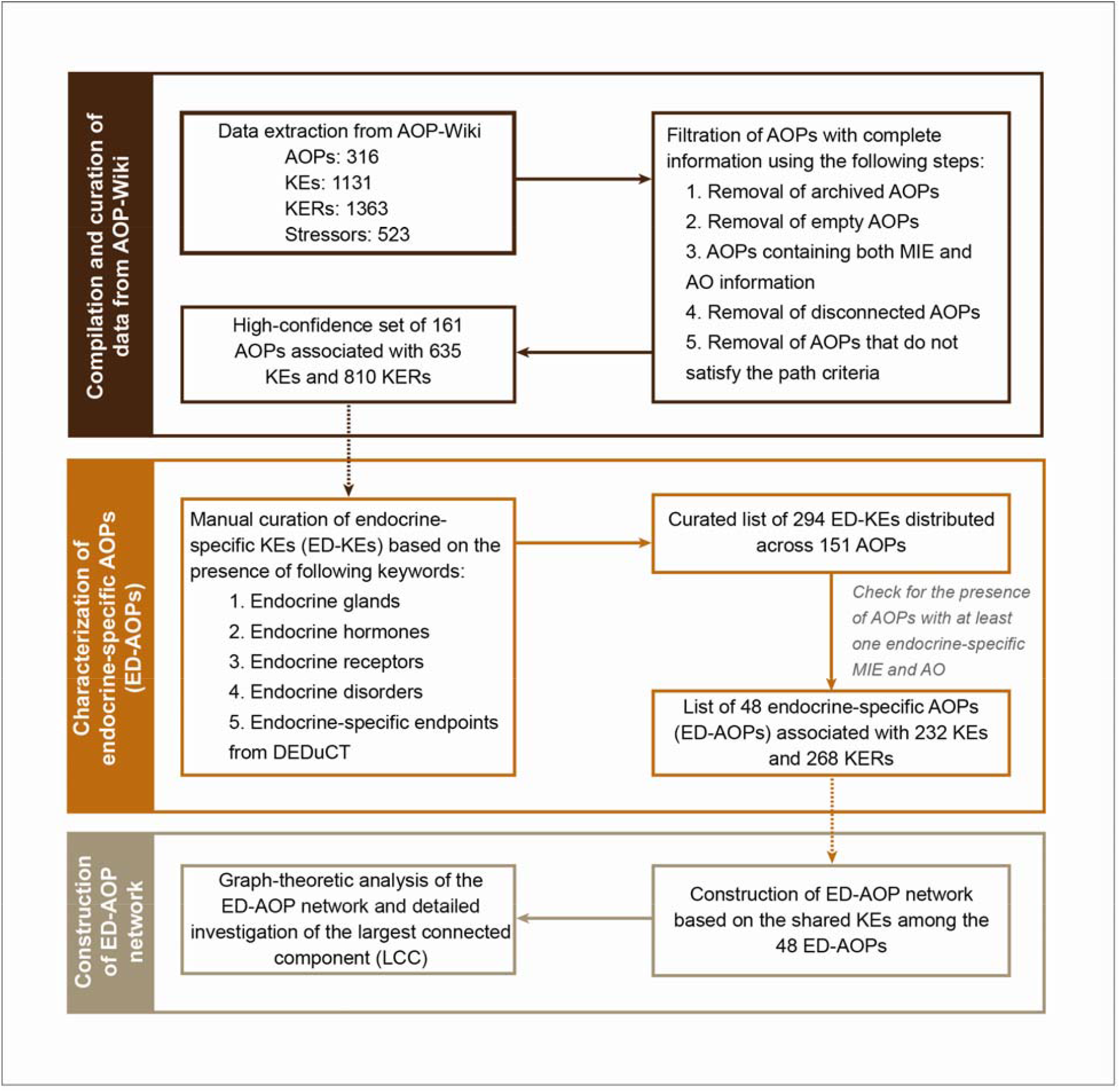
Detailed workflow for the development, characterization and analysis of an adverse outcome pathway (AOP) network for endocrine disruption.

## 2. Methods

### 2.1. Compilation of AOP dataset from AOP-Wiki

From the ‘Project Downloads’ (https://aopwiki.org/downloads) section of the AOP-Wiki, we have downloaded the XML archive as on 03 January 2021. This XML archive from AOP-Wiki was parsed using the xml2 package in R to obtain information on 316 AOPs, 1131 Key Events (KEs), 1363 Key-Event Relationships (KERs), and 523 stressors.

For each AOP in AOP-Wiki, we have retrieved information including the AOP identifier, AOP title, OECD status, and Society for the Advancement of AOPs (SAAOP) status. For each KE in an AOP, we have gathered information including the KE identifier, KE type, level of biological organization and taxonomy. The KE type can be either molecular initiating event (MIE), key event (KE) or adverse outcome (AO). For each KER in an AOP, we have gathered information including the KER identifier, upstream KE, downstream KE, the weight of evidence (WoE), adjacency information, and the quantitative understanding score. Lastly, we have compiled the chemical stressors linked to KEs in different AOPs along with their structure information such as the CAS identifier (https://www/cas.ohttps://www.epa.gov/chemical-research/distributed-structure-searchable-toxicity-dsstox-database) and InChIKey. Note that the AOP-Wiki also contains information on non-chemical stressors such as genetic or environmental factors.

We remark that each AOP can be viewed as a directed graph or network wherein the nodes are KEs and directed edges are KERs linking upstream KEs with downstream KEs. In this directed graph representation of an AOP, it is straightforward to determine the existence of a directed path between any pair of KEs.

### 2.2. Filtration of high-confidence AOPs from AOP-Wiki

Since AOP-Wiki is under continuous development, some AOPs may have incomplete information (OECD, 2018). Therefore, it is important to evaluate the quality and completeness of information in each AOP before their selection for the derived AOP network construction (Villeneuve et al., 2018a). We have assessed the quality and completeness of information in each AOP obtained from AOP-Wiki as follows (Figure 1).

Firstly, we have removed the ‘archived AOPs’ based on SAAOP status as these are no longer under active development. This led to the removal of 6 AOPs. Secondly, we have removed ‘empty AOPs’ which do not have a KE or a KER. After removing ‘archived AOPs’ and ‘empty AOPs’, we have 218 AOPs that remain under consideration. Thirdly, we have removed any AOP which does not contain at least one MIE and at least one AO. After this step, we have 182 AOPs with both MIE and AO that remain under consideration. Fourthly, we have computed the number of (weakly) connected components in each AOP. This led to the identification of 3 disconnected AOPs that have more than one connected component. After the removal of 3 disconnected AOPs, we have 179 AOPs that remain under consideration.

Fifthly, we have computed directed paths from different MIEs to different AOs in each AOP to filter out incomplete AOPs. Since an AOP can have both multiple MIEs and multiple AOs, we have computed the directed paths between each pair of MIE and AO in an AOP to impose this path criterion. We have retained an AOP only if it satisfies the following path criteria:

a. Every MIE in an AOP has at least one (outgoing) path to at least one AO in the same AOP.
b. Every AO in an AOP has at least one (incoming) path from at least one MIE in the same AOP.
c. Every KE in an AOP (other than MIEs and AOs) has at least one incoming path from at least one MIE in the same AOP and at least one outgoing path to at least one AO in the same AOP. After removing AOPs that do not satisfy the path criteria, we arrive at a high-confidence set of 161 AOPs which are associated with 635 KEs and 810 KERs (Figure 1; Supplementary Table S1). Next, these 161 high-confidence AOPs were considered for the identification of AOPs relevant for endocrine disruption.

### 2.3. Curated subset of endocrine-specific AOPs

To build the AOP network specific to endocrine disruption, it is important to identify the subset of endocrine-specific AOPs (ED-AOPs) among the 161 high-confidence AOPs. To identify ED-AOPs, we have manually curated the endocrine-specific KEs (ED-KEs) among the 635 KEs associated with the 161 high-confidence AOPs.

A KE was identified as an ED-KE if the KE contains keywords relevant to the endocrine system. Keywords relevant to endocrine system were identified based on: (a) List of endocrine glands, (b) List of endocrine hormones, (c) List of endocrine receptors where hormones can bind, (d) List of endocrine disorders in MeSH (https://meshb.nlm.nih.gov/), and (e) List of endocrine-specific endpoints in DEDuCT (Karthikeyan et al., 2021, 2019). This process led to a curated subset of 294 ED-KEs (Supplementary Table S2). Afterwards, we retained 151 AOPs among the 161 high-confidence AOPs that contain at least one ED-KE. Furthermore, we consider an AOP to be an ED-AOP if it contains at least one MIE which is an ED-KE and at least one AO which is an ED-KE. This filtration led to a curated subset of 48 ED-AOPs which are associated with 232 KEs and 268 KERs (Table 1; Figure 1; Supplementary Table S3).

**Table 1:** The curated subset of 48 ED-AOPs among the 161 high-confidence AOPs filtered from AOP-Wiki. The table also gives the fraction of ED-KEs, the cumulative WoE score, and the WoE score for human applicability (Human WoE) for each of the 48 ED-AOPs.

| Serial number | AOP identifier | AOP title | Fraction of ED-KEs (%) | Cumulative WoE | Human WoE |
| --- | --- | --- | --- | --- | --- |
| 1 | 6 | Antagonist binding to PPARα leading to body-weight loss | 62.5 | High | High |
| 2 | 7 | Aromatase (Cyp19a1) reduction leading to impaired fertility in adult female | 100 | High | Low |
| 3 | 12 | Chronic binding of antagonist to N-methyl-D-aspartate receptors (NMDARs) during brain development leads to neurodegeneration with impairment in learning and memory in aging | 87.5 | Moderate | Low |
| 4 | 13 | Chronic binding of antagonist to N-methyl-D-aspartate receptors (NMDARs) during brain development induces impairment of learning and memory abilities | 90 | Low | High |
| 5 | 18 | PPARα activation in utero leading to impaired fertility in males | 87.5 | Moderate | Low |
| 6 | 19 | Androgen receptor antagonism leading to adverse effects in the male foetus (mammals) | 100 | - | - |
| 7 | 28 | Cyclooxygenase inhibition leading reproductive failure | 66.7 | Moderate | - |
| 8 | 36 | Peroxisomal Fatty Acid Beta-Oxidation Inhibition Leading to Steatosis | 75 | High | - |
| 9 | 37 | PPARα activation leading to hepatocellular adenomas and carcinomas in rodents | 80 | High | - |
| 10 | 41 | Sustained AhR Activation leading to Rodent Liver Tumours | 100 | High | - |
| 11 | 42 | Inhibition of Thyroperoxidase and Subsequent Adverse Neurodevelopmental Outcomes in Mammals | 62.5 | High | High |
| 12 | 43 | Disruption of VEGFR Signaling Leading to Developmental Defects | 60 | High | Moderate |
| 13 | 60 | NR1I2 (Pregnane X Receptor, PXR) activation leading to hepatic steatosis | 58.3 | High | - |
| 14 | 62 | AKT2 activation leading to hepatic steatosis | 100 | - | - |
| 15 | 63 | Cyclooxygenase inhibition leading to reproductive dysfunction | 80 | Moderate | Low |
| 16 | 64 | Glucocorticoid Receptor (GR) Mediated Adult Leydig Cell Dysfunction Leading to Decreased Male Fertility | 100 | - | - |
| 17 | 100 | Cyclooxygenase inhibition leading to reproductive dysfunction via inhibition of female spawning behavior | 57.1 | Moderate | - |
| 18 | 101 | Cyclooxygenase inhibition leading to reproductive dysfunction via inhibition of pheromone release | 57.1 | High | - |
| 19 | 102 | Cyclooxygenase inhibition leading to reproductive dysfunction via interference with meiotic prophase I /metaphase I transition | 80 | High | Low |
| 20 | 103 | Cyclooxygenase inhibition leading to reproductive dysfunction via interference with spindle assembly checkpoint | 80 | High | Low |
| 21 | 107 | Constitutive androstane receptor activation leading to hepatocellular adenomas and carcinomas in the mouse and the rat | 80 | High | - |
| 22 | 110 | Inhibition of iodide pump activity leading to follicular cell adenomas and carcinomas (in rat and mouse) | 100 | - | - |
| 23 | 111 | Decrease in androgen receptor activity leading to Leydig cell tumors (in rat) | 100 | - | - |
| 24 | 112 | Increased dopaminergic activity leading to endometrial adenocarcinomas (in Wistar rat) | 100 | - | - |
| 25 | 117 | Androgen receptor activation leading to hepatocellular adenomas and carcinomas (in mouse and rat) | 75 | - | - |
| 26 | 118 | Chronic cytotoxicity leading to hepatocellular adenomas and carcinomas (in mouse and rat) | 75 | - | - |
| 27 | 119 | Inhibition of thyroid peroxidase leading to follicular cell adenomas and carcinomas (in rat and mouse) | 100 | - | - |
| 28 | 120 | Inhibition of 5α-reductase leading to Leydig cell tumors (in rat) | 100 | - | - |
| 29 | 124 | HMG-CoA reductase inhibition leading to decreased fertility | 83.3 | - | - |
| 30 | 164 | Beta-2 adrenergic agonist activity leading to mesovarian leiomyomas in the rat and mouse | 66.7 | High | - |
| 31 | 167 | Early-life estrogen receptor activity leading to endometrial carcinoma in the mouse. | 71.4 | High | - |
| 32 | 201 | Juvenile hormone receptor agonism leading to male offspring induction associated population decline | 50 | - | - |
| 33 | 203 | 5-hydroxytryptamine transporter inhibition leading to decreased reproductive success and population decline | 37.5 | - | - |
| 34 | 204 | 5-hydroxytryptamine transporter inhibition leading to increased reproductive success and population increase | 37.5 | - | - |
| 35 | 205 | AOP from chemical insult to cell death | 50 | High | - |
| 36 | 212 | Histone deacetylase inhibition leading to testicular atrophy | 66.7 | Moderate | Moderate |
| 37 | 216 | Excessive reactive oxygen species production leading to population decline via follicular atresia | 85.7 | - | - |
| 38 | 220 | Cyp2E1 Activation Leading to Liver Cancer | 80 | High | Moderate |
| 39 | 232 | NFE2/Nrf2 repression to steatosis | 87.5 | - | - |
| 40 | 271 | Inhibition of thyroid peroxidase leading to impaired fertility in fish | 80 | High | - |
| 41 | 293 | Increased DNA damage leading to increased risk of breast cancer | 66.7 | Moderate | - |
| 42 | 299 | Excessive reactive oxygen species production leading to population decline via reduced fatty acid beta-oxidation | 62.5 | - | - |
| 43 | 300 | Thyroid Receptor Antagonism and Subsequent Adverse Neurodevelopmental Outcomes in Mammals | 40 | Moderate | High |
| 44 | 306 | Androgen receptor (AR) antagonism leading to short anogenital distance (AGD) in male (mammalian) offspring | 100 | High | Moderate |
| 45 | 336 | DNA methyltransferase inhibition leading to population decline (1) | 57.1 | Moderate | - |
| 46 | 337 | DNA methyltransferase inhibition leading to population decline (2) | 62.5 | Moderate | - |
| 47 | 340 | DNA methyltransferase inhibition leading to transgenerational effects (1) | 62.5 | Moderate | - |
| 48 | 341 | DNA methyltransferase inhibition leading to transgenerational effects (2) | 66.7 | Moderate | - |

Subsequently, we have studied the enrichment of ED-KEs across these 48 ED-AOPs by computing the fraction of ED-KEs among KEs in an ED-AOP. Among the curated subset of 48 ED-AOPs, we find that 11 ED-AOPs are such that 100% (all) of their KEs are ED-KEs, and moreover, 45 ED-AOPs are such that at least 50% of their KEs are ED-KEs. Note that the minimum fraction of ED-KEs in an ED-AOP among the 48 ED-AOPs is found to be 37.5% (Table 1).

Furthermore, we have computed a cumulative weight of evidence (cumulative WoE) score for each of the 48 ED-AOPs based on the weight of evidence (WoE) scores given by AOP-Wiki to the associated 268 KERs. For each KER, the AOP-Wiki gives one of the following values namely, ‘high’, ‘moderate’, ‘low’ or ‘not specified’ as the WoE score, and this value is a measure of the strength of empirical evidence supporting the casual relationship between the pair of KEs connected by a KER. Note that the WoE scores assigned to KERs by AOP-Wiki can change with updates in the resource (OECD, 2018; Pollesch et al., 2019). Also, different KERs in any AOP can differ in their WoE scores. Therefore, we propose the following cumulative WoE score for each ED-AOP based on the WoE scores given by AOP-Wiki to associated KERs.

For each ED-AOP, we compute the fraction of KERs with different values of the WoE score namely, ‘high’, ‘moderate’, ‘low’ or ‘not specified’. For example, the fraction of KERs in an ED-AOP with WoE score ‘high’ can be computed from the ratio of the number of KERs in the AOP with WoE score ‘high’ and the total number of KERs in the AOP, and this quantity for an ED-AOP is denoted by F(‘high’). Similarly, it is straightforward to compute the quantities F(‘moderate’), F(‘low’) and F(‘not specified’) for an ED-AOP. In a nutshell, for each of the 48 ED-AOPs, we have computed the quantities F(‘high’), F(‘moderate’), F(‘low’) and F(‘not specified’) from the WoE scores of the associated KERs (Supplementary Table S4). Subsequently, we have assigned the cumulative WoE score to each ED-AOP as follows:

i. If an ED-AOP has F(‘high’) ≥ 0.5, then the cumulative WoE score was assigned to ‘high’.
ii. Else if an ED-AOP has F(‘high’) < 0.5 but has [F(‘high’) + F(‘moderate’)] ≥ 0.5, then the cumulative WoE score was assigned to ‘moderate’.
iii. Else if an ED-AOP has [F(‘high’) + F(‘moderate’)] < 0.5 but has [F(‘high’) + F(‘moderate’) + F(‘low’)] ≥ 0.5, then the cumulative WoE score was assigned to ‘low’.
iv. Else if an ED-AOP has [F(‘high’) + F(‘moderate’) + F(‘low’)] < 0.5, then the cumulative WoE score was assigned to ‘not specified’.

Based on this definition, we find that 18, 12, 1 and 17 ED-AOPs were assigned cumulative WoE score of ‘high’, ‘moderate’, ‘low’ and ‘not specified’, respectively (Table 1; Supplementary Table S4).

In Supplementary Table S5, we compile the biological domain information namely, taxonomic, sex and life stage applicability, for each ED-AOP from AOP-Wiki. For example, AOP:13 (https://aopwiki.org/aops/13) is ‘Chronic binding of antagonist to N-methyl-D-aspartate receptors (NMDARs) during brain development induces impairment of learning and memory abilities’. The taxonomic applicability information for AOP:13 indicates that the AOP is applicable to human, mouse, rat and monkey. The sex applicability information for AOP:13 indicates that the AOP is applicable to both sexes (male and female). The life stage applicability information for AOP:13 indicates that the AOP is relevant during brain development (Supplementary Table S5). Similar to WoE scores for KERs in AOP-Wiki, the WoE information for taxonomic, sex, or life stage applicability for each AOP in AOP-WiKi can have one of the four values namely, ‘high’, ‘moderate’, ‘low’ or ‘not specified’. Lastly, we have evaluated the information on taxonomic applicability of the 48 ED-AOPs from AOP-Wiki to assess the human applicability of each ED-AOP. We find that 14 out of the 48 ED-AOPs have evidence for human applicability in AOP-Wiki (Table 1; Supplementary Table S5). Of these 14 ED-AOPs with evidence for human applicability, 4, 4 and 6 ED-AOPs have WoE score for human applicability to be ‘high’, ‘moderate’ and ‘low’, respectively (Table 1; Supplementary Table S5). Note that if the WoE score for taxonomic applicability of an ED-AOP for *Homo sapiens* was ‘not specified’ in AOP-Wiki, we have assigned the WoE score for human applicability of that ED-AOP in Table 1 to ‘low’.

Evidently, the cumulative WoE score and the WoE score for human applicability listed in Table 1 can be used to assess the level of evidence for an ED-AOP and further filter the curated subset of 48 ED-AOPs. Nevertheless, we have not imposed any filters based on taxonomic, sex, or life stage applicability information in AOP-Wiki during the filtration of the 48 ED-AOPs for the subsequent construction of the derived AOP network.

### 2.4. Construction of the ED-AOP network and its connected components

After filtration of the curated subset of 48 ED-AOPs, we have constructed the AOP network specific to endocrine disruption by assembling the information on shared KEs and KERs among the 48 ED-AOPs. We refer to this derived AOP network as ‘ED-AOP network’ (Figure 2). The ED-AOP network contains KEs and KERs across the 48 ED-AOPs, and thus, captures diverse biological perturbations related to endocrine system (Knapen et al., 2018; Villeneuve et al., 2018a). The ED-AOP network can be visualized as an undirected graph of 48 nodes corresponding to the 48 ED-AOPs, and there exists an edge between any two nodes in this undirected graph if the two ED-AOPs have at least one shared KE (Figure 2).

**Figure 2:**
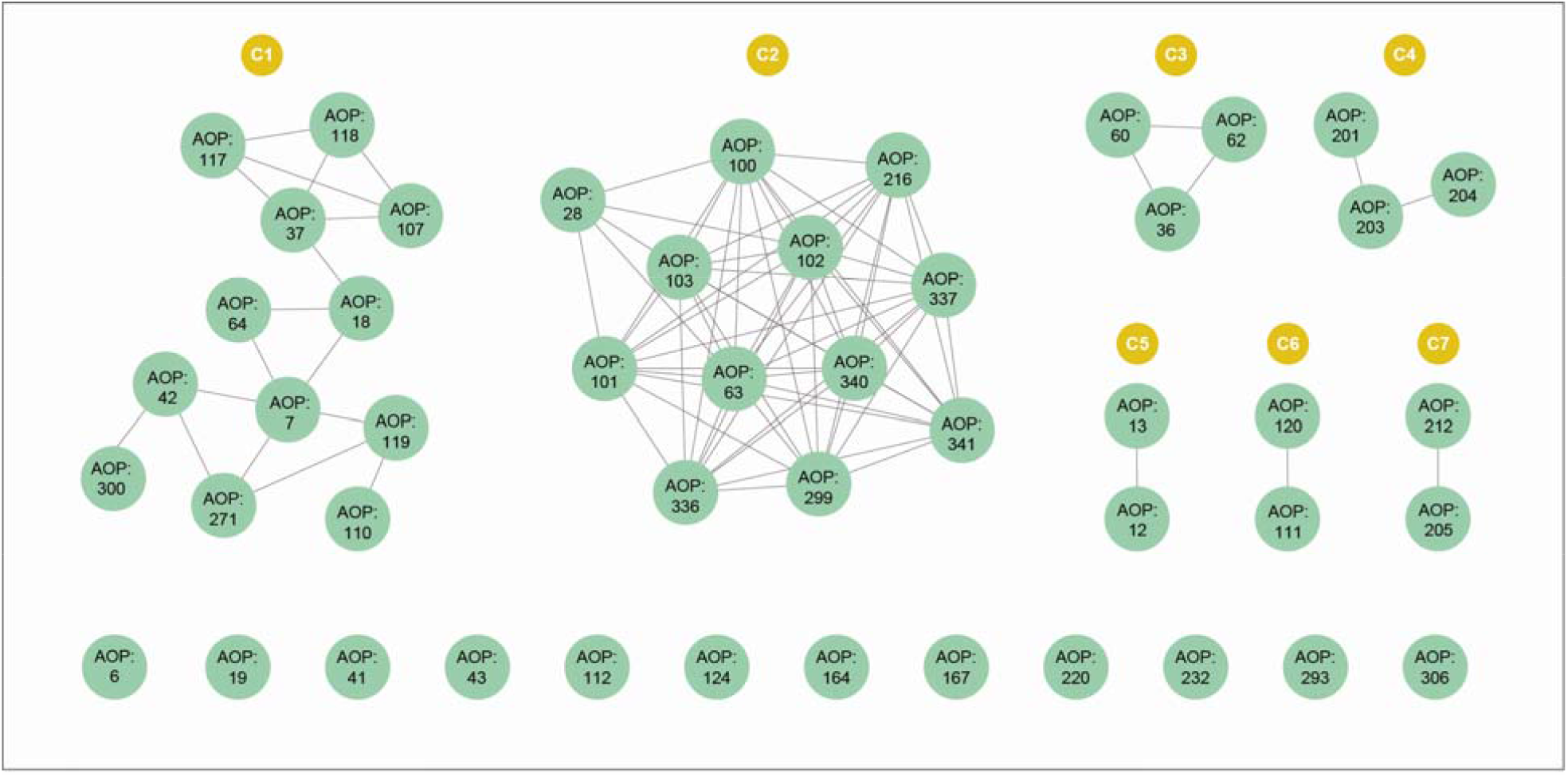
Visualization of the ED-AOP network based on shared KEs among the 48 ED-AOPs. Here, each node corresponds to an ED-AOP and there exists an edge between any two ED-AOPs if they have at least one shared KE. The network has 7 connected components (labeled C1-C7) with ≥ 2 ED-AOPs and 12 isolated ED-AOPs. The two largest connected components (LCCs) labeled by C1 and C2 contain 12 ED-AOPs each.

In this study, we have performed a graph-theoretic analysis of the ED-AOP network to reveal important topological features (Villeneuve et al., 2018a). To assess the overall connectivity of the ED-AOP network, we have computed the connected components using python package NetworkX (https://networkx.org/). A connected component is a subset of nodes in the graph wherein there exists at least one path between every pair of nodes in the induced subgraph. Note that a completely connected network has a single connected component comprising all nodes in the graph. Based on this computation, we find that the ED-AOP network can be decomposed into 7 connected components with ≥ 2 ED-AOPs and 12 isolated ED-AOPs. These 7 connected components together comprise 36 ED-AOPs (Figure 2; Supplementary Table S6). Among these 7 connected components, the two largest connected components (LCCs) labeled by C1 and C2 in Figure 2 contain 12 ED-AOPs each, and the remaining 5 connected components contain ≤ 3 ED-AOPs each.

### 2.5. Detailed analysis of the largest connected components of the ED-AOP network

In this study, we have performed a detailed analysis of the two LCCs, C1 and C2, in the ED-AOP network. For this purpose, we have constructed the directed network corresponding to each LCC wherein nodes are KEs and edges represent KERs in the 12 ED-AOPs comprising the LCC of the ED-AOP network (Figures 3 and 4). Subsequently, we have studied four standard network measures namely, in-degree, out-degree, betweenness centrality and eccentricity, for KEs in the directed network corresponding to LCC, and these measures were computed using NetworkAnalyzer (https://apps.cytoscape.org/apps/networkanalyzer) in Cytoscape (Assenov et al., 2008). In the directed network, in-degree (respectively, out-degree) of a KE refers to the number of KEs immediately upstream (respectively, immediately downstream) of that KE (Villeneuve et al., 2018a). Importantly, in-degree and out-degree of KEs can help identify points of convergence and divergence in the directed network. Further, betweenness centrality can help identify KEs crucial for the spread of biological perturbations, while eccentricity can help identify KEs which are farthest upstream or farthest downstream in the directed network (Spinu et al., 2019; Villeneuve et al., 2018a). By applying network measures, we have studied the systems-level perturbations caused by endocrine-mediated events in the ED-AOP network upon chemical exposure. We have also investigated the ED-AOP network for possible emergence of new paths between pairs of MIE and AO that are both ED-KEs and belong to different ED-AOPs.

**Figure 3:**
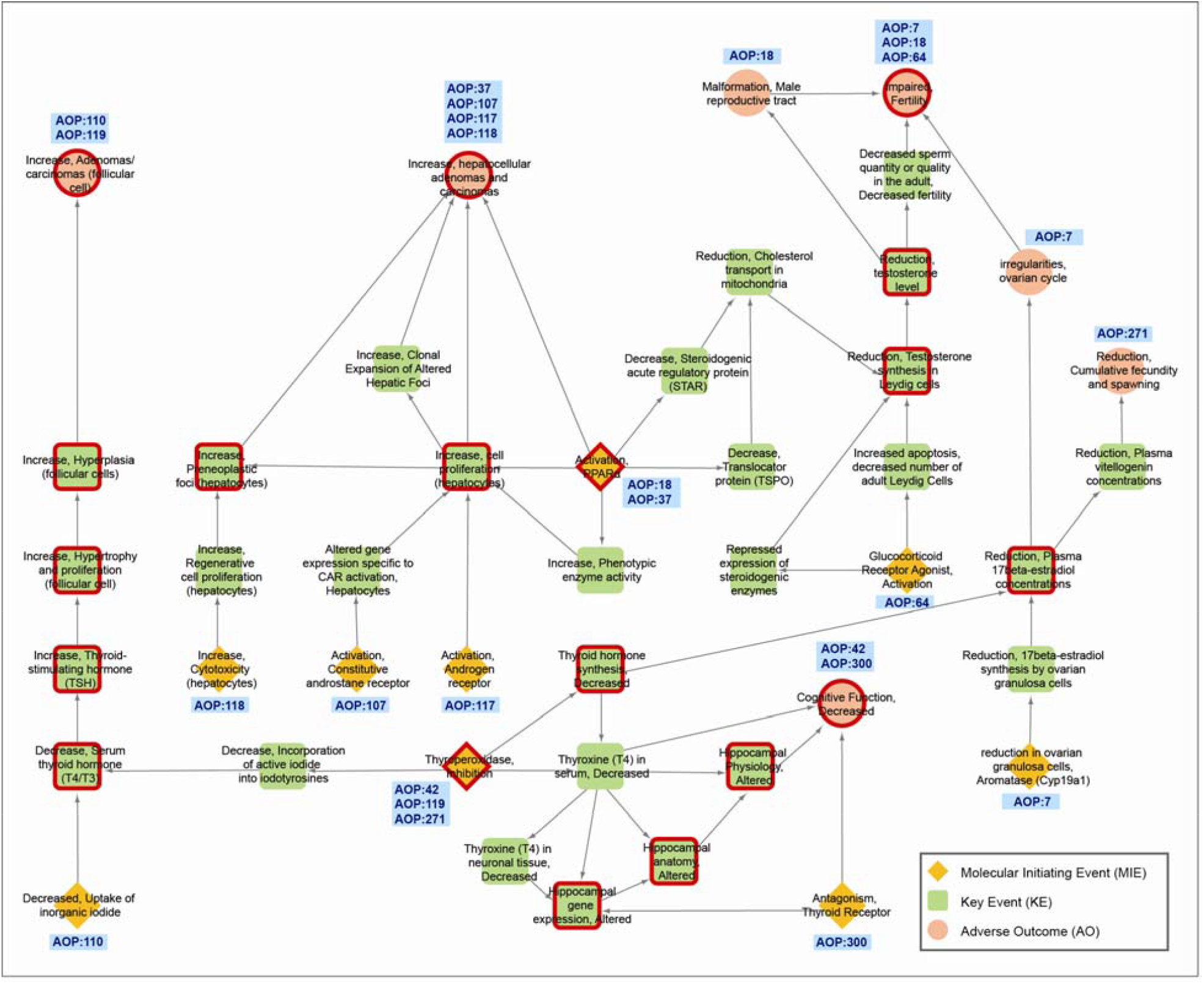
The directed network for LCC C1 in the ED-AOP network consisting of 44 KEs and 56 KERs. The 44 KEs in C1 can be categorized into 9 MIEs, 28 KEs and 7 AOs. MIEs, KEs and AOs are shown in distinct shapes namely, diamond, square and circle, respectively. The 19 shared KEs in C1 are marked in ‘red’. For each MIE and AO, the corresponding AOP identifier is displayed in this figure.

**Figure 4:**
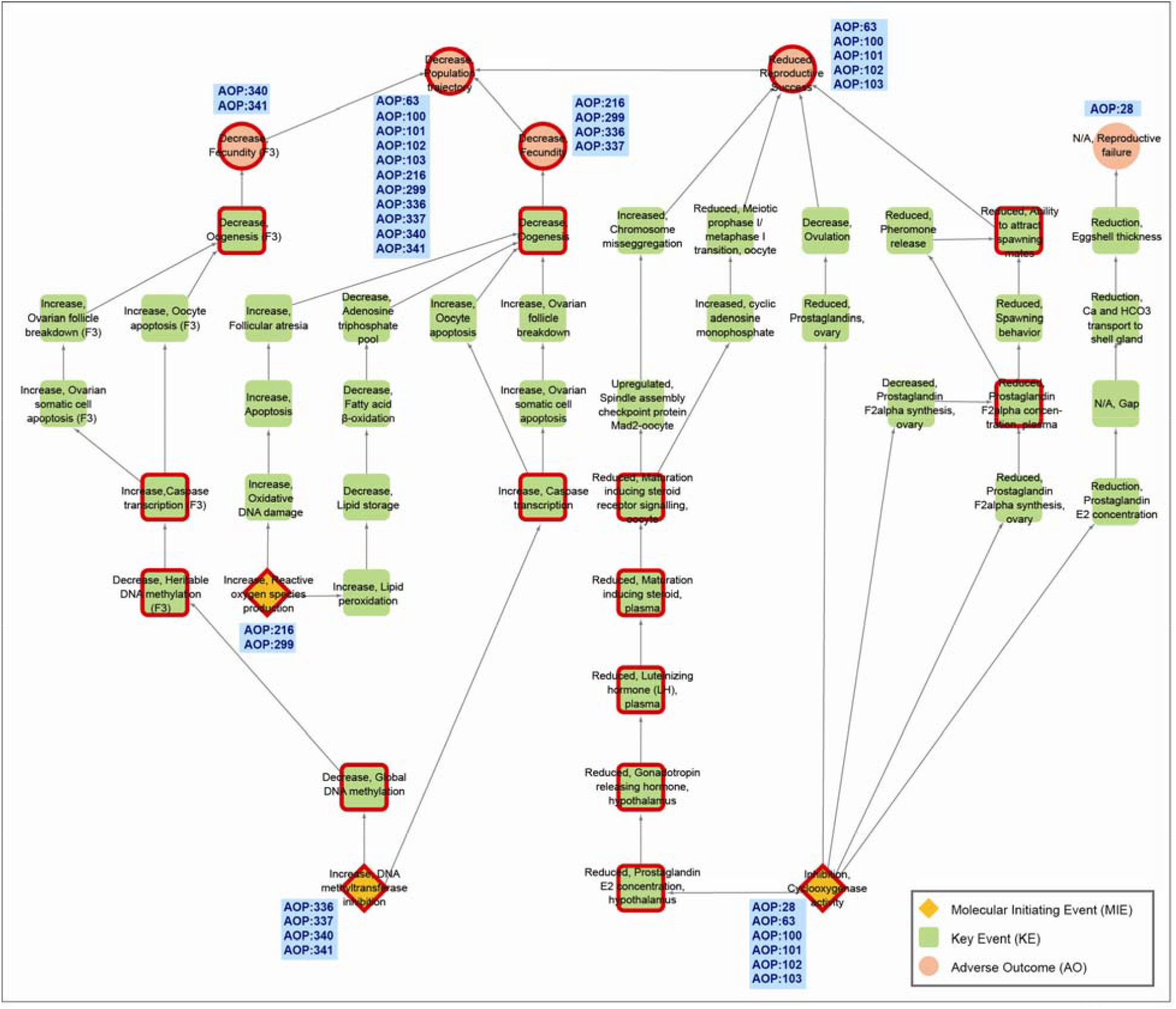
The directed network for LCC C2 in the ED-AOP network consisting of 48 KEs and 56 KERs. The 48 KEs in C2 can be categorized into 3 MIEs, 40 KEs and 5 AOs. MIEs, KEs and AOs are shown in distinct shapes namely, diamond, square and circle, respectively. The 20 shared KEs in C2 are marked in ‘red’. For each MIE and AO, the corresponding AOP identifier is displayed in this figure.

Finally, we have analyzed the information from AOP-Wiki on stressors associated with KEs in an ED-AOP network. The stressors with incomplete information in AOP-Wiki were manually assigned to their structural identifiers including CAS, DSSTOX and InChIKey. Based on this exercise, we identified 35 and 16 chemical stressors associated with KEs in the two LCCs, C1 and C2, respectively. Thereafter, the list of 792 EDCs in DEDuCT 2.0 (Karthikeyan et al., 2021, 2019) was used to identify the subset of known EDCs among the chemical stressors associated with C1 and C2. Finally, we identified 16 and 6 EDCs associated with components C1 and C2, respectively.

## 3. Results and Discussion

### 3.1. Derived AOP network specific to endocrine disruption

The aim of this study is to develop a derived AOP network specific to endocrine disruption based on information in AOP-Wiki. To construct this ‘ED-AOP network’, we have compiled detailed information on 316 AOPs, 1131 KEs and 1363 KERs from AOP-Wiki (Methods). Due to continuous development of AOP-Wiki, some AOPs may have incomplete information at any particular time. Therefore, we have filtered a high-confidence subset of 161 AOPs among the 316 AOPs in AOP-Wiki as follows (Methods; Figure 1).

Firstly, we excluded 6 ‘archived AOPs’ that are not under active development. Secondly, we excluded the ‘empty AOPs’ that contain neither a KE nor a KER, and this led to a subset of 218 AOPs. Thirdly, we retained AOPs that contain both MIE and AO, and this led to a subset of 182 AOPs. Fourthly, we excluded AOPs with two or more connected components, and this led to a subset of 179 AOPs. Fifthly, we retained AOPs that satisfy the path criteria as described in the Methods section. Each of the 179 AOPs retained in the previous step satisfy the criteria whereby: (i) there is at least one outgoing path from every MIE to at least one AO in the AOP, and (ii) there is at least one incoming path to every AO from at least one MIE in the AOP. However, 18 out of the 179 AOPs retained in the previous step fail to satisfy the criteria whereby every KE (other than a MIE or an AO) has at least one incoming path from at least one MIE in the AOP and has at least one outgoing path to at least one AO in the AOP. Exclusion of these 18 AOPs that fail the path criteria results in the subset of 161 high-confidence AOPs (Figure 1; Supplementary Table S1).

Afterwards, we identified ED-AOPs among the 161 high-confidence AOPs as follows (Methods; Figure 1). Among the 635 KEs associated with these 161 AOPs, we manually filtered a subset of 294 ED-KEs (Methods; Supplementary Table S2). We next retained 151 out of the 161 high-confidence AOPs that contain at least one ED-KE. Further, we retained 48 out of these 151 AOPs that contain at least one MIE and at least one AO which are ED-KEs, and these 48 AOPs were designated as ED-AOPs (Methods; Table 1; Supplementary Table S3).

An analysis of the fraction of ED-KEs in each ED-AOP finds that this fraction varies in the range 37.5% to 100% across the 48 ED-AOPs. Specifically, 11 ED-AOPs have 100% (all) of their KEs as ED-KEs, while 45 ED-AOPs have at least 50% of their KEs as ED-KEs (Table 1). Moreover, we propose a cumulative WoE score for each ED-AOP based on the WoE scores assigned to KERs by AOP-Wiki (Methods). Based on our definition, 18, 12, 1 and 17 ED-AOPs were assigned cumulative WoE scores of ‘high’, ‘moderate’, ‘low’ and ‘not specified’, respectively (Table 1). Further, we have assessed the information from AOP-Wiki on taxonomic, sex, or life stage applicability of the 48 ED-AOPs (Methods). We find that 14 out of these 48 ED-AOPs have evidence for human applicability, of which 4, 4 and 6 ED-AOPs can be assigned WoE scores for human applicability of ‘high’, ‘moderate’ and ‘low’, respectively (Table 1).

Thereafter, we have constructed the ED-AOP network by leveraging the information on shared KEs among the 48 ED-AOPs (Methods). Figure 2 visualizes this ED-AOP network as an undirected graph wherein nodes correspond to 48 ED-AOPs and edges between pairs of ED-AOPs signify that they have at least one KE in common. Subsequently, we investigated the overall connectivity of this ED-AOP network by determining the connected components in this undirected graph (Methods). It can be seen that the ED-AOP network can be divided into 7 connected components with ≥ 2 ED-AOPs (which are labeled C1 to C7 in Figure 2) and 12 isolated ED-AOPs. Simply stated, 36 ED-AOPs within these 7 connected components have at least one KE which is shared with at least one other ED-AOP belonging to the same connected component, while the remaining 12 isolated ED-AOPs do not have any shared KE. Importantly, there are two LCCs, labeled C1 and C2 in Figure 2, of 12 ED-AOPs each, among the 7 connected components in the ED-AOP network. The LCCs C1 and C2 comprise of 44 and 48 KEs, respectively, of which 19 and 20 KEs are shared among 2 or more ED-AOPs in C1 and C2, respectively (Figures 3 and 4).

To better understand the systems-level effects of AOs in the 7 components of the ED-AOP network, we have categorized AOs into 4 systems-level endocrine-mediated perturbations, namely, ‘hepatic’, ‘metabolic’, ‘neurological’ and ‘reproductive’, and this classification depends on the perturbed biological process corresponding to an AO (Table 2). For example, the AO titled ‘Increase, hepatocellular adenomas and carcinomas’ in AOP-Wiki was classified as ‘hepatic’ while the AO titled ‘impaired, Fertility’ as ‘reproductive’ (Table 2). This categorization of AOs in ED-AOPs into 4 systems-level perturbations follows a similar classification scheme for observed adverse effects upon exposure to endocrine disrupting chemicals (EDCs) in our previous work (Karthikeyan et al., 2021, 2019). We observe that majority of AOs in the ED-AOP network affect the ‘reproductive’ system (Table 2). Moreover, the AOs in C1 can affect 4 different systems, while all AOs in C2 affect solely the ‘reproductive’ system (Table 2).

**Table 2:** The list of AOs in the 7 connected components of the ED-AOP network and their categorization into 4 systems-level endocrine-mediated perturbations, namely, ‘hepatic’, ‘metabolic’, ‘neurological’ and ‘reproductive’, depending on the perturbed biological processes.

| Serial number | Component identifier | AO | System-level perturbation |
| --- | --- | --- | --- |
| 1 | C1 | Increase, hepatocellular adenomas and carcinomas | Hepatic |
| 2 | C1 | Increase, Adenomas/carcinomas (follicular cell) | Metabolic |
| 3 | C1 | Cognitive Function, Decreased | Neurological |
| 4 | C1 | impaired, Fertility | Reproductive |
| 5 | C1 | irregularities, ovarian cycle | Reproductive |
| 6 | C1 | Reduction, Cumulative fecundity and spawning | Reproductive |
| 7 | C1 | Malformation, Male reproductive tract | Reproductive |
| 8 | C2 | Decrease, Population trajectory | Reproductive |
| 9 | C2 | Decrease, Fecundity | Reproductive |
| 10 | C2 | Decrease, Fecundity (F3) | Reproductive |
| 11 | C2 | N/A, Reproductive failure | Reproductive |
| 12 | C3 | Increased, Liver Steatosis | Hepatic |
| 13 | C4 | Increased, Male offspring | Reproductive |
| 14 | C4 | Decreased, Reproductive Success | Reproductive |
| 15 | C4 | Increased, Reproductive Success | Reproductive |
| 16 | C5 | Impairment, Learning and memory | Neurological |
| 17 | C6 | Increase, Leydig cell tumors | Reproductive |
| 18 | C7 | Apoptosis | - |
| 19 | C7 | Testicular atrophy | Reproductive |

### 3.2. Topological analysis of the largest components in the ED-AOP network

Since the two LCCs dominate the ED-AOP network, we decided to next focus on them. For a detailed analysis of each LCC in the ED-AOP network, we have constructed the corresponding directed network wherein nodes are KEs and each directed edge represents a KER linking its upstream KE with its downstream KE (Methods). In Figures 3 and 4, we display the directed networks corresponding to LCCs C1 and C2, respectively, wherein nodes are KEs and directed edges are KERs. The directed network for C1 (Figure 3) has 44 KEs and 56 KERs while that for C2 (Figure 4) has 48 KEs and 56 KERs. To identify important events (KEs) leading to endocrine disruption in the ED-AOP network, we have next computed node-centric measures namely, in-degree, out-degree, betweenness centrality and eccentricity, for KEs in the directed networks for C1 and C2.

Firstly, we identified convergent and divergent events within the directed networks for C1 and C2 by assessing the in-degree and out-degree of each KE. A KE is considered to be ‘convergent’ if the in-degree is greater than (>) out-degree for the particular KE, while a KE is considered to be ‘divergent’ if the in-degree is less than (<) out-degree for the particular KE (Villeneuve et al., 2018a). In C1, there are 13 convergent KEs and 12 divergent KEs. Among the 13 convergent KEs in C1, 2 KEs namely, ‘Increase, cell proliferation (hepatocytes)’ and ‘Increase, hepatocellular adenomas and carcinomas’, have the highest in-degree of 4. Among the 12 divergent KEs in C1, 2 KEs namely, ‘Activation, PPARα’ and ‘Thyroxine (T4) in serum, Decreased’, have the highest out-degree of 5, and in other words, these 2 divergent events lead to 5 other events in C1 (Figure 3; Supplementary Table S7). In C2, there are 6 convergent KEs and 7 divergent KEs. Among the 6 convergent KEs in C2, 2 KEs namely, ‘Decrease, Oogenesis’ and ‘Reduced, Reproductive Success’, have the highest in-degree of 4. Among the 7 divergent KEs in C2, the KE ‘Inhibition, Cyclooxygenase activity’ has the highest out-degree of 5 (Figure 4; Supplementary Table S7).

Secondly, we have assessed the betweenness centrality of KEs in the directed networks for C1 and C2. The shared KE ‘Reduction, Testosterone synthesis in Leydig cells’ has the maximum betweenness centrality of 0.4 in C1 (Supplementary Figure S1; Supplementary Table S7), while the shared KE ‘Reduced, Maturation inducing steroid receptor signalling, oocyte’ has the maximum betweenness centrality of 0.43 in C2 (Supplementary Figure S2; Supplementary Table S7). Since these KEs with the highest betweenness centrality are on the shortest paths linking various nodes in C1 or C2, the events serve as significant control points in the ED-AOP network (Leydesdorff, 2007).

Thirdly, we have assessed the eccentricity of KEs in the directed networks for C1 and C2. The higher the eccentricity value for a node, the farther is the node located with respect to other nodes in the network, and thus, low eccentricity value for a node indicates its central location in the network (Takes and Kosters, 2011). In C1, the 2 shared KEs namely, ‘Activation, PPARα’ and ‘Thyroperoxidase, Inhibition’, have the maximum eccentricity value of 6 (Supplementary Figure S3; Supplementary Table S7). In C2, the shared KE ‘Reduced, Prostaglandin E2 concentration, hypothalamus’ has the maximum eccentricity value of 8 (Supplementary Figure S4; Supplementary Table S7).

Afterwards, we assessed the available information in AOP-Wiki for the two LCCs, C1 and C2. For C1, 21 out of the 44 KEs, i.e. nearly 50%, have evidence for human applicability in AOP-Wiki. For C2, however, 46 out of the 48 KEs do not have taxonomic applicability information in AOP-Wiki. Further, C2 contains two pairs of ED-AOPs namely, (i) AOP:336 and AOP:337, and (ii) AOP:340 and AOP:341, such that each pair of ED-AOPs contain the identical set of MIEs and AOs (Supplementary Table S6). Further, each pair of ED-AOPs is such that the two ED-AOPs have most of their KEs in common, and thus, it may be worthwhile to consider only one ED-AOP in each pair to avoid duplication of information in the ED-AOP network. Moreover, we find that AOP:28 of C2 contains KEs such as ‘N/A, Gap’ and ‘N/A, Reproductive failure’. Overall, this highlights disparity and gaps in available information across AOPs in AOP-Wiki. In sum, the available information is more comprehensive for the 12 ED-AOPs in C1 (in comparison to C2). As a result, the LCC C1 was further investigated to reveal the systems-level perturbations caused by endocrine-mediated events, the emergence of new paths linking MIEs and AOs, and the chemical stressors associated with KEs.

### 3.3. Systems-level perturbations caused by endocrine-mediated events in the largest component C1 of the ED-AOP network

Human exposure to EDCs can lead to endocrine disruption that in turn can affect various biological systems. Of late, there is concern regarding an increase in the incidence of endocrine-mediated disorders linked to reproduction, metabolism, development, nervous system and immunity in humans and wildlife (Bergman et al., 2013; Diamanti-Kandarakis et al., 2009; Gore et al., 2015; Zoeller et al., 2012). To better understand the systems-level perturbations upon EDC exposure, it is important to investigate the associated endocrine-mediated events leading to varied adverse outcomes. In this direction, we have investigated the systems-level perturbations caused by endocrine-mediated events captured in LCC C1 of the ED-AOP network.

In LCC C1, there are 44 KEs of which 9 are MIEs and 7 are AOs. Notably, 37 out of these 44 KEs (84%) in C1 were found to be ED-KEs. Depending on the perturbed cell types, organs or biological processes, we categorized the 44 KEs in C1 into 4 different systems-level endocrine-mediated perturbations, namely, ‘hepatic’, ‘metabolic’, ‘neurological’ and ‘reproductive’ (Figure 5; Supplementary Table S8). This categorization scheme for the 44 KEs in C1 is similar to the one used for AOs listed in Table 2. For example, the KE titled ‘Increase, Phenotypic enzyme activity’ (https://aopwiki.org/events/1170) in AOP-Wiki is associated with the cellular term ‘hepatocyte’, and thus, the KE is categorized as ‘hepatic’ in our scheme (Figure 5; Supplementary Table S8). However, the information on the perturbed cell types, organs or biological processes is not available in AOP-Wiki for 3 MIEs in C1, namely, ‘Antagonism, Thyroid Receptor’, ‘Activation, Androgen receptor’, and ‘Activation, Constitutive androstane receptor’, and this prevented the categorization of these 3 MIEs into any of the 4 different systems-level perturbations (Figure 5; Supplementary Table S8). Of the remaining 41 KEs in C1, 9, 10, 5, and 17 KEs were categorized as ‘hepatic’, ‘metabolic’, ‘neurological’ and ‘reproductive’ systems-level perturbations, respectively (Figure 5; Supplementary Table S8).

**Figure 5:**
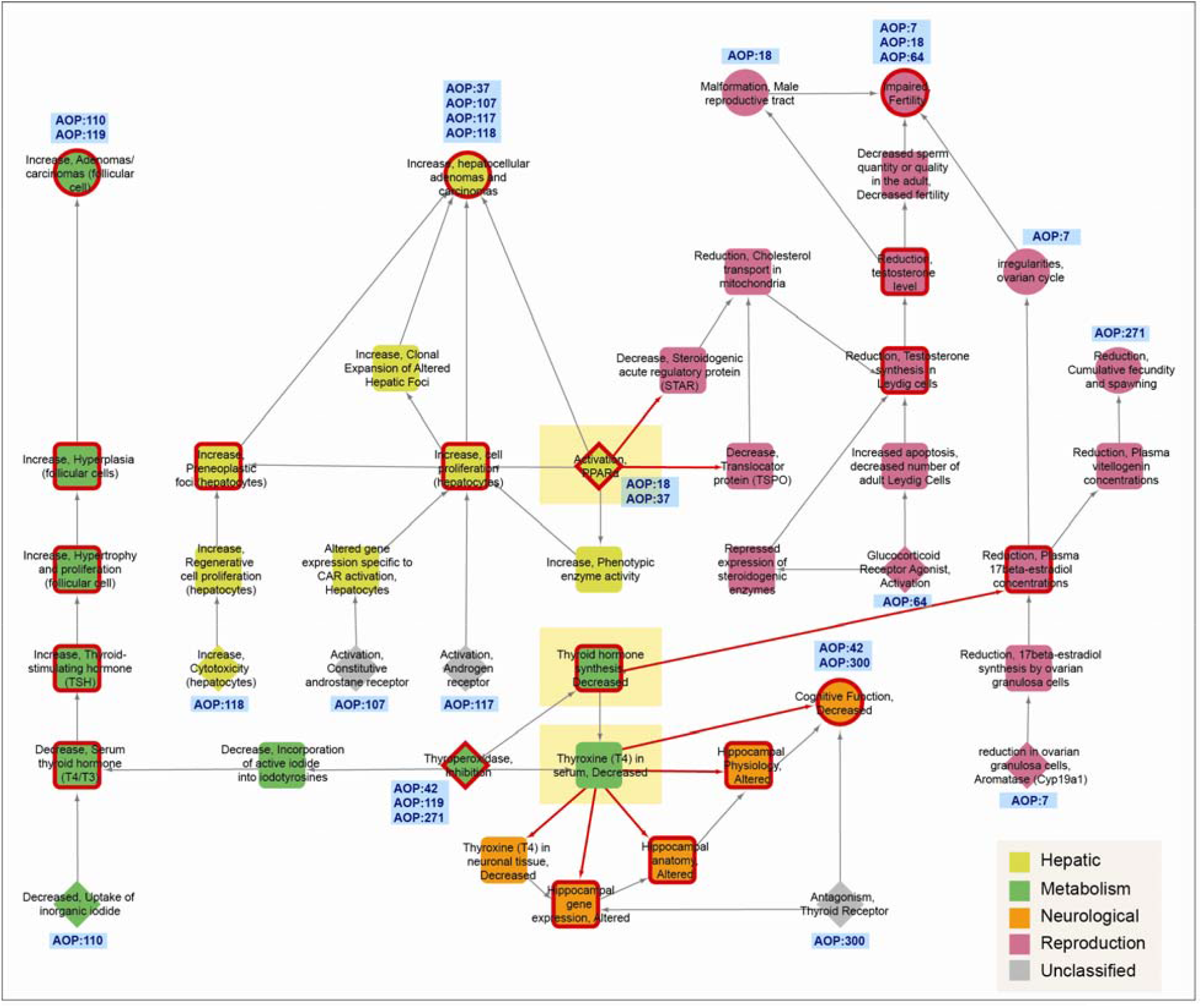
The directed network for LCC C1 in the ED-AOP network consisting of 44 KEs wherein the KEs are colored based on their categorization into 4 systems-level perturbations namely, hepatic, metabolic, neurological and reproductive. MIEs, KEs and AOs are shown in distinct shapes namely, diamond, square and circle, respectively. The 19 shared KEs in C1 are marked in ‘red’. The ‘red’ edges highlight KERs that connect KEs categorized into different systems-level perturbations. The ‘yellow rectangles’ highlight 3 divergent KEs which serve as *point of divergence* from one system to another system.

Thereafter, we analyzed the topology of LCC C1 by considering the categorization of the 41 KEs into 4 different systems-level perturbations (Figure 5). Specifically, we determined KERs in C1 that connect two KEs that differ in their categorization into systems-level perturbations. We find 8 such KERs in C1 of which 5 KERs connect KEs in metabolic and neurological systems, 2 KERs connect KEs in hepatic and reproductive systems, and 1 KER connects KEs in metabolic and reproductive systems (Figure 5). Among the KEs associated with these 8 KERs, 3 (divergent) KEs namely, ‘Activation, PPARα’, ‘Thyroxine (T4) in serum, Decreased’, and ‘Thyroid hormone synthesis, Decreased’, serve as *points of divergence* linking different systems in C1 (Figure 5). Specifically, the divergent KE titled ‘Thyroxine (T4) in serum, Decreased’ is categorized as ‘metabolic’ by our scheme, and this KE is immediately upstream of 5 KEs namely, ‘Thyroxine (T4) in neuronal tissue, Decreased’, ‘Hippocampal gene expression, Altered’, ‘Hippocampal anatomy, Altered’, ‘Hippocampal Physiology, Altered’, and ‘Cognitive Function, Decreased’, categorized as ‘neurological’ in C1 (Figure 5). In other words, this analysis of C1 reveals that the metabolic event ‘Thyroxine (T4) in serum, Decreased’ can lead to 5 neurological events, and interestingly, we were able to find independent supporting evidence for these particular associations between metabolic and neurological events in the published literature (Bernal, 2000; Cooke et al., 2014; Tunc-Ozcan et al., 2013; Volpato et al., 2002).

Furthermore, the divergent KE titled ‘Activation, PPARα’ is categorized as ‘hepatic’, and this KE is immediately upstream of 2 KEs namely, ‘Decrease, Steroidogenic acute regulatory protein (STAR)’ and ‘Decrease, Translocator protein (TSPO)’, categorized as ‘reproductive’ in C1 (Figure 5), and there are supporting evidences for these particular associations between hepatic and reproductive events in the published literature (Batarseh and Papadopoulos, 2010; Corton and Lapinskas, 2005; Latini et al., 2008). Finally, the divergent KE titled ‘Thyroid hormone production, Decreased’ is categorized as ‘metabolic’, and this KE is immediately upstream of a KE titled ‘Reduction, Plasma 17beta-estradiol concentrations’ categorized as ‘reproductive’ in C1 (Figure 5), and there is supporting evidence for this particular association on the influence of thyroid levels on reproductive hormones (Saran et al., 2016). Thus, analysis of divergent KEs in the ED-AOP network can offer insights into links between different systems affected by endocrine disruption.

Lastly, we observed that 4 out of 12 ED-AOPs in C1 contain a shared AO titled ‘Increase, hepatocellular adenomas and carcinomas’ which is categorized as ‘hepatic’ systems-level perturbation. In C1, the shared KE titled ‘Increase, cell proliferation (hepatocytes)’ has maximum in-degree and is an important *point of convergence* leading to the above-mentioned AO. Further, this convergent KE is downstream of MIEs linked to activation of three hormonal receptors namely, Constitutive Androstane receptor (CAR), Androgen receptor (AR), and PPARα, highlighting the possibility of additive or synergistic effects upon exposure to EDCs targeting multiple receptors (Figure 5). In sum, the analysis of divergent or convergent KEs in the ED-AOP network from the perspective of associated systems-level perturbations can aid in the risk assessment of endocrine disruptors.

### 3.4. Emergent paths in the ED-AOP network

Since an AOP network contains multiple AOPs connected via shared KEs, new (directed) paths, other than those in individual AOPs, can emerge between MIEs and AOs belonging to different AOPs in the corresponding directed network of KEs and KERs (Methods). Such emergent paths from MIEs to AOs in an AOP network can also lead to the development of new stand-alone AOPs (Villeneuve et al., 2018a). Here, we have investigated the possibility of such emergent paths between MIEs and AOs in the LCC C1 of the ED-AOP network consisting of 12 ED-AOPs (Methods). We have found 4 new paths in the LCC C1 that connect an endocrine-specific MIE in one ED-AOP to an endocrine-specific AO in another ED-AOP (Figure 3; Table 3).

**Table 3:** The table gives information on the starting MIE and the ending AO for each of the 4 new paths identified in the LCC C1 of the ED-AOP network.

| Serial number | MIE | AO |
| --- | --- | --- |
| 1 | Thyroperoxidase, Inhibition | irregularities, ovarian cycle |
| 2 | Thyroperoxidase, Inhibition | impaired, Fertility |
| 3 | reduction in ovarian granulosa cells, Aromatase (Cyp19a1) | Reduction, Cumulative fecundity and spawning |
| 4 | Glucocorticoid Receptor Agonist, Activation | Malformation, Male reproductive tract |

Of the 4 new paths in C1 (Figure 3; Table 3), 2 new paths start from the shared MIE ‘Thyroperoxidase, Inhibition’ (in AOP:42, AOP:119, and AOP:271) and end at the 2 AOs namely, ‘irregularities, ovarian cycle’ (in AOP:7) and ‘impaired, Fertility’ (in AOP:7, AOP:18, and AOP:64). Another new path in C1 starts from the MIE ‘reduction in ovarian granulosa cells, Aromatase (Cyp19a1)’ (in AOP:7) and ends at the AO ‘Reduction, Cumulative fecundity and spawning’ (in AOP:271). Lastly, there is a new path in C1 starting from the MIE ‘Glucocorticoid Receptor Agonist, Activation’ (in AOP:64) and ending at the AO ‘Malformation, Male reproductive tract’ (in AOP:18). These emergent paths identified in LCC C1 of the ED-AOP network have potential to reveal unknown relationships between distant KEs and may represent toxicity pathways specific to endocrine disruption. Further, a closer inspection of these emergent paths may also lead to prediction of unknown adverse effects upon specific EDC exposure.

### 3.5. Chemical stressors and the ED-AOP network

Chemical risk assessment is the primary objective for the development of AOPs (Ankley et al., 2010; Villeneuve et al., 2014a). For proper risk assessment, it is important to link chemical exposure to perturbations in biological systems or processes (Ankley et al., 2010). Thus, we have analyzed the information on chemical stressors associated with KEs in LCC C1 from AOP-Wiki (Methods). Based on information in AOP-Wiki, 35 chemical stressors were found to be associated with different KEs in C1. By performing a comparative analysis of these 35 chemical stressors with the list of 792 potential EDCs in DEDuCT 2.0 (Karthikeyan et al., 2021, 2019), we identified a subset of 16 chemical stressors associated with C1 that have strong supporting evidence of endocrine disruption (Methods; Supplementary Table S9). These 16 EDCs directly target at least one event among 5 MIEs, 9 KEs and 2AOs in LCC C1. Among these 5 MIEs, MIE ‘Thyroperoxidase, Inhibition’ is directly linked to 7 EDCs, and MIE ‘Activation, PPARα’ is directly linked to 4 EDCs. Among the 16 EDCs, we find that the EDC ‘6-Propyl-2-thiouracil’ directly targets 8 events in C1 (Supplementary Table S9).

Analyses of direct associations between chemical stressors and KEs in the ED-AOP network can reveal the diversity of biological mechanisms via which EDCs can cause different endocrine-mediated adverse effects. To aid ongoing efforts in risk assessment of EDCs, it will be worthwhile to undertake a future effort to associate all known EDCs, including 792 potential EDCs in DEDuCT 2.0, to different events in the ED-AOP network. However, such an effort is beyond the scope of this work. In sum, a stressor-ED-AOP network can serve as a predictive model for EDCs and their adverse effects.

## 4. Conclusions

An AOP is a systematic framework to encapsulate the existing toxicological information as a toxicity pathway to aid in risk assessment and chemical regulation (Ankley et al., 2010; Edwards et al., 2016; Vinken, 2018, 2013; Vinken et al., 2017). Within AOP-Wiki, the up to date central repository of individual AOPs, AOP networks have emerged due to sharing of KEs and KERs across individual AOPs. Since AOP networks are expected to be the functional units for prediction in real-world scenarios, there is notable interest in the derivation and analysis of AOP networks tailored to address specific problems or applications (Knapen et al., 2018; Pollesch et al., 2019; Villeneuve et al., 2018a).

The challenges in the risk assessment and regulation of EDCs partially stem from the existing knowledge gaps in linking chemical exposure to diverse adverse outcomes (Bergman et al., 2013; Darbre, 2015; Fuhrman et al., 2015). To address this challenge, a blueprint of the endocrine disruption mechanisms in the form of toxicity pathways spanning different levels of biological organization can be invaluable (Browne et al., 2020). In this context, the development of a comprehensive AOP network specific to endocrine disruption (i.e., an ED-AOP network) can aid ongoing research and policy framing surrounding EDCs. In this work, we have developed a detailed workflow (Figure 1) to leverage information in AOP-Wiki and construct a comprehensive ED-AOP network (Figure 2; Table 1). Ensuing graph-theoretic analysis of this ED-AOP network of 48 ED-AOPs, and in particular, its largest components C1 and C2 of 12 ED-AOPs each, reveals several mechanistic insights on endocrine-mediated perturbations upon chemical exposure.

Since AOP development is a continuous and iterative exercise, therefore the ED-AOP network constructed in this study is limited by the existing knowledge in AOP-Wiki. As AOPs are living documents, it will be important to maintain the ED-AOP network up to date with any expansion in AOP-Wiki. We expect that the detailed workflow in Figure 1 with a little or no modification can be used for any future update of the ED-AOP network. Moreover, the current information in AOP-Wiki on chemical stressors associated with events in the ED-AOP network is a small fraction of the existing knowledge on potential EDCs in the published literature (see Results and Discussion) (Karthikeyan et al., 2021, 2019), and therefore, it will be important to invest future efforts towards developing a comprehensive stressor-ED-AOP network wherein all known EDCs are linked to different events in the ED-AOP network. In sum, the ED-AOP network along with information on chemical stressors will enable better risk assessment and regulation of EDCs.

## Author contribution statement

**Janani Ravichandran:** Conceptualization, Data curation, Formal analysis, Software, Visualization, Writing – original draft. **Bagavathy Shanmugam Karthikeyan:** Formal analysis, Writing – original draft. **Areejit Samal:** Conceptualization, Supervision, Formal analysis, Writing – original draft.

## Acknowledgements

Areejit Samal is supported by the Science and Engineering Research Board (SERB) India (through a Ramanujan fellowship (SB/S2/RJN-006/2014)) and the Max Planck Society Germany (through a Max Planck Partner Group in Mathematical Biology). The funding agencies have no role in study design, data collection, data analysis, manuscript preparation or decision to publish.

## Supplementary Figures

**Figure S1:**
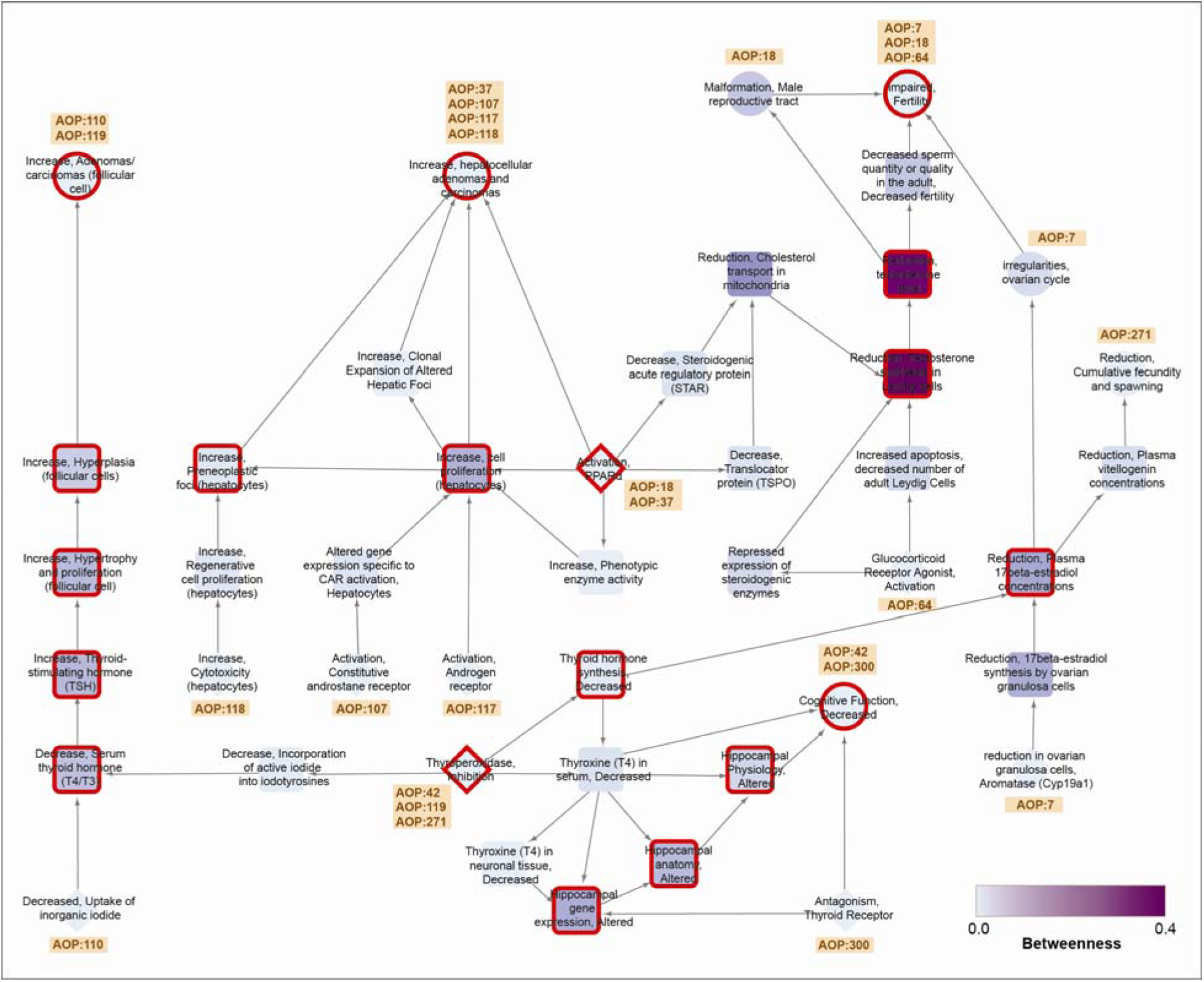
The directed network for LCC C1 wherein the KEs are colored based on their betweenness centrality values. MIEs, KEs and AOs are shown in distinct shapes namely, diamond, square and circle, respectively. The shared KEs are marked in ‘red’. For each MIE and AO, the AOP identifier is displayed in this figure.

**Figure S2:**
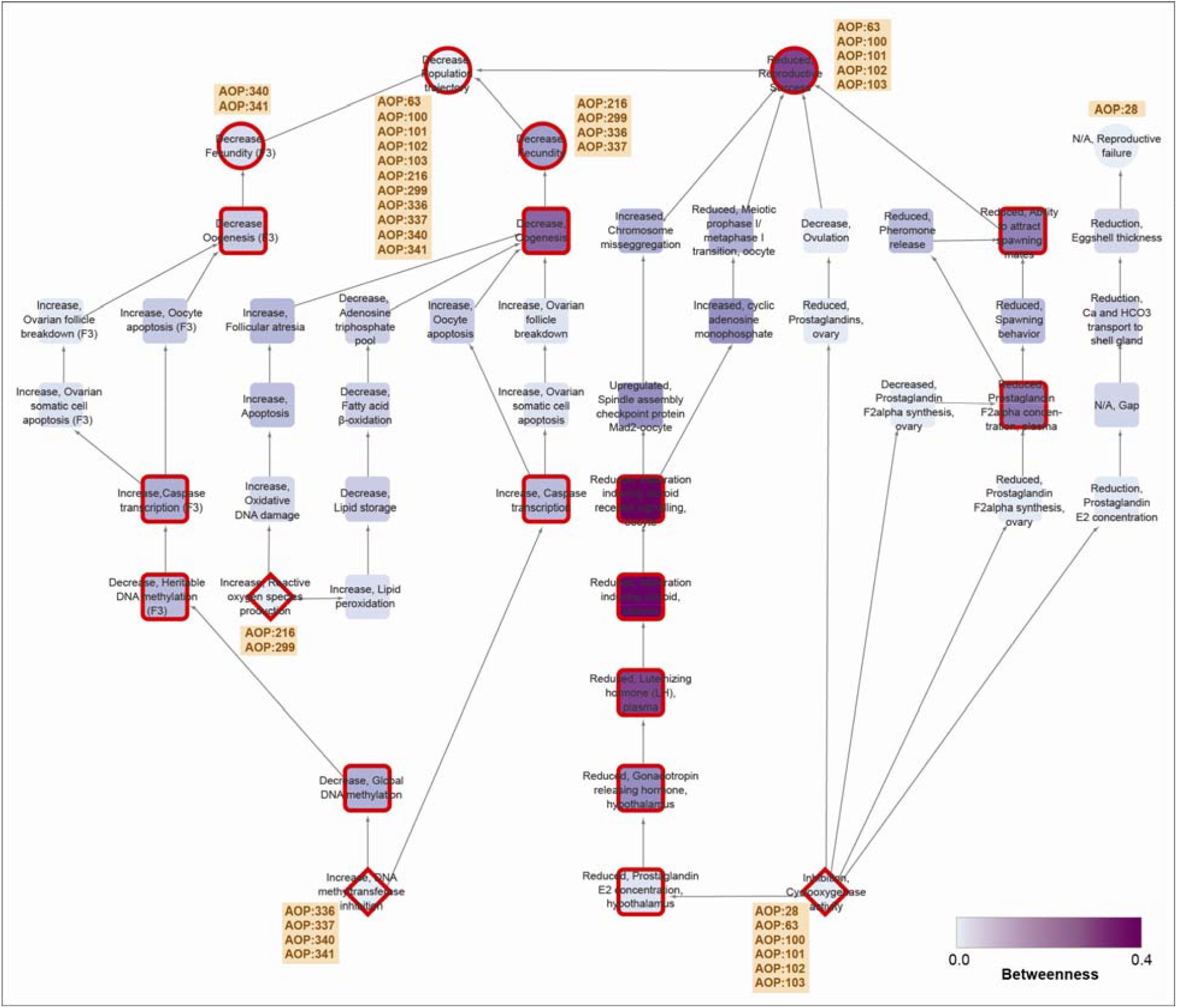
The directed network for LCC C2 wherein the KEs are colored based on their betweenness centrality values. MIEs, KEs and AOs are shown in distinct shapes namely, diamond, square and circle, respectively. The shared KEs are marked in ‘red’. For each MIE and AO, the AOP identifier is displayed in this figure.

**Figure S3:**
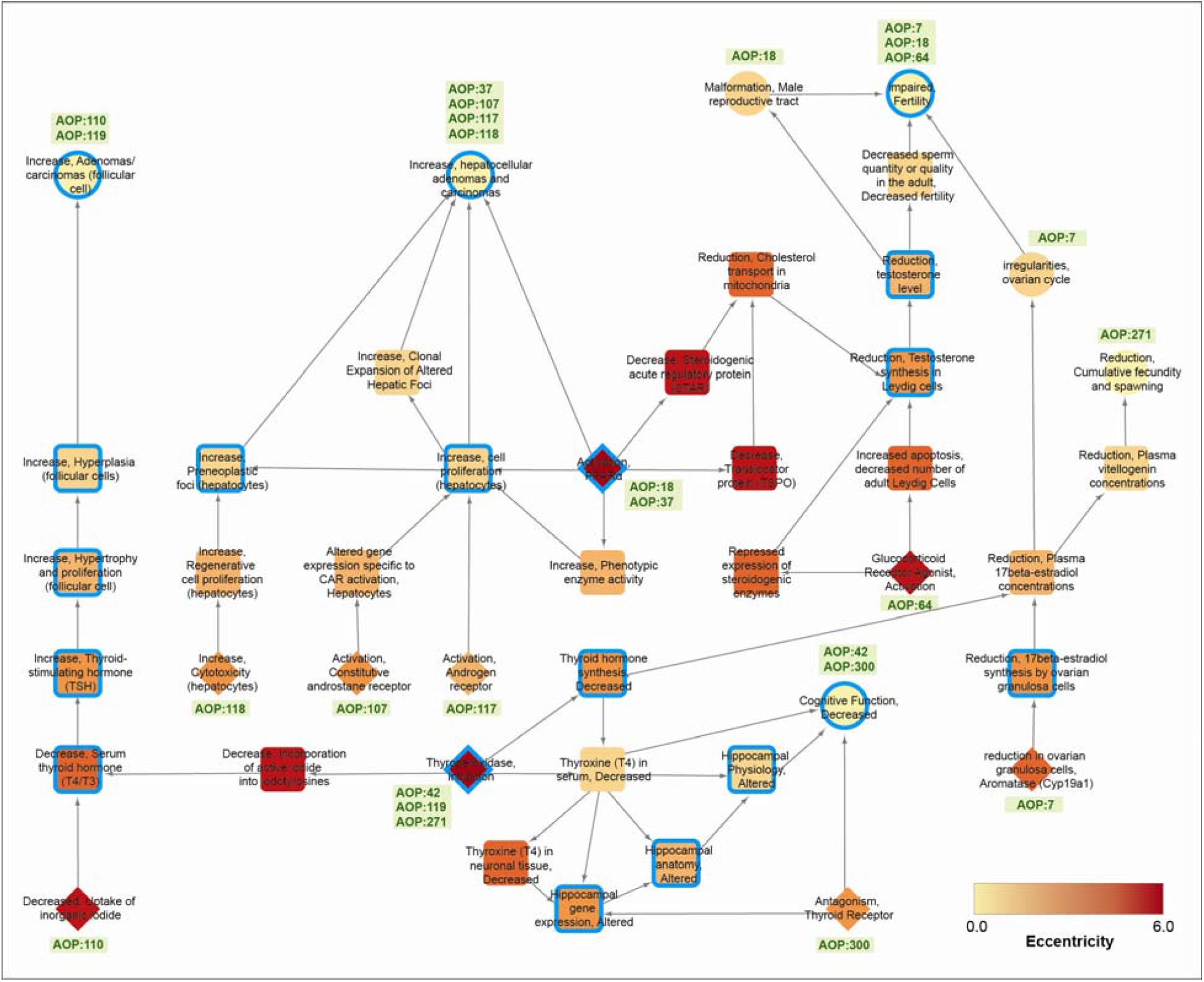
The directed network for LCC C1 wherein the KEs are colored based on their eccentricity values. MIEs, KEs and AOs are shown in distinct shapes namely, diamond, square and circle, respectively. The shared KEs are marked in ‘red’. For each MIE and AO, the AOP identifier is displayed in this figure.

**Figure S4:**
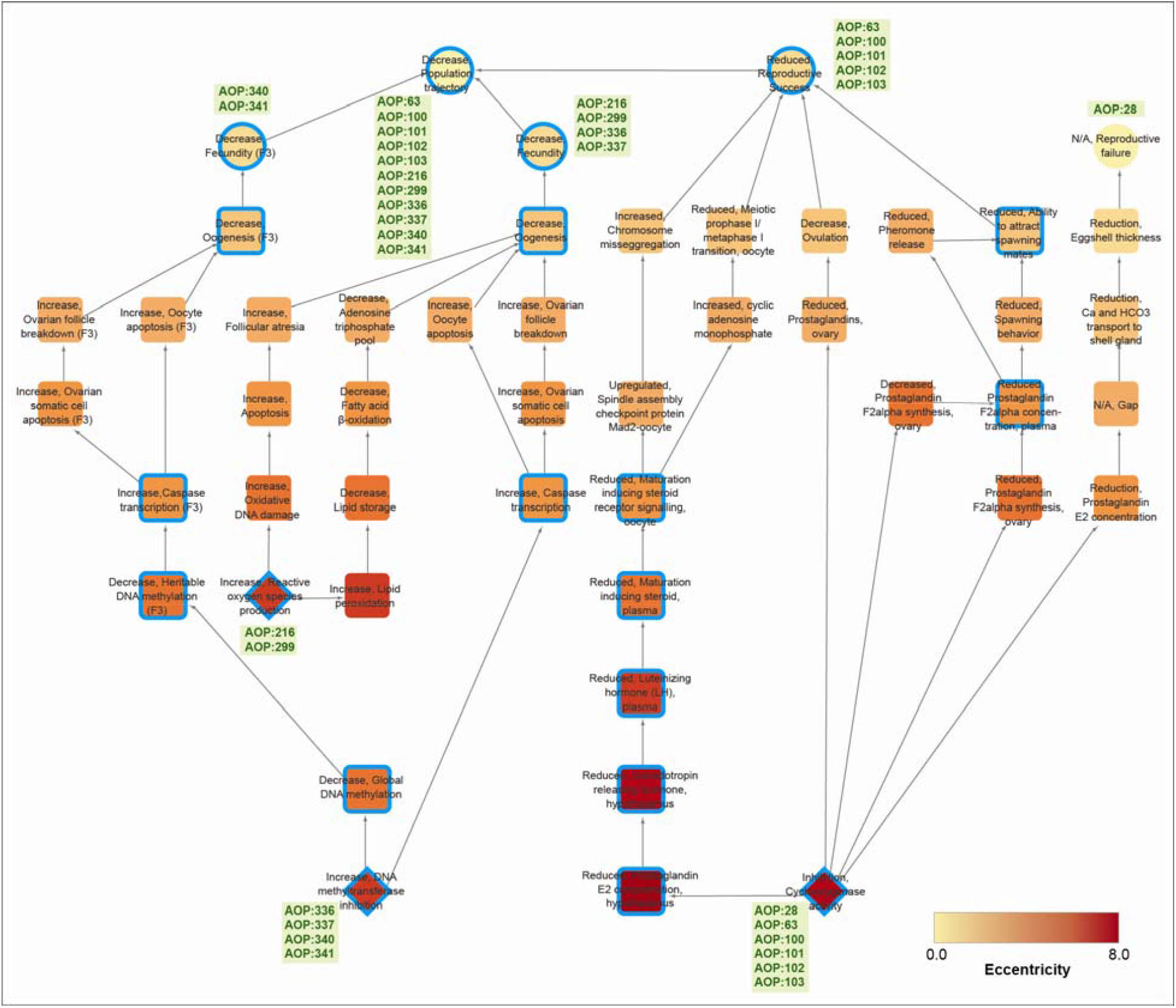
The directed network for LCC C2 wherein the KEs are colored based on their eccentricity values. MIEs, KEs and AOs are shown in distinct shapes namely, diamond, square and circle, respectively. The shared KEs are marked in ‘red’. For each MIE and AO, the AOP identifier is displayed in this figure.

## Captions of Supplementary Tables

**Table S1:** Filtered list of 161 high-confidence AOPs with complete information from AOP-Wiki.

**Table S2:** Manually curated list of 294 endocrine-specific key events (ED-KEs) among the 635 KEs associated with the 161 high-confidence AOPs.

**Table S3:** The curated subset of 48 ED-AOPs along with their associated KEs and KERs. For each KER, the table gives the KER identifier, upstream event, downstream event, the weight of evidence (WoE), adjacency information, and the quantitative understanding score.

**Table S4:** The table gives the fraction of KERs with each value of WoE score, namely ‘high’, ‘moderate’, ‘low’ or ‘not specified’, for the 48 ED-AOPs. The table also gives the cumulative WoE score for each ED-AOP.

**Table S5:** The table provides additional layers of information associated with the 48 ED-AOPs including Taxonomic Applicability, Sex Applicability, and Life stage Applicability from AOP-Wiki along with the corresponding weight of evidence (WoE) scores. The column ‘Human WoE’ gives the weight of evidence score for human applicability for each of the 48 ED-AOPs.

**Table S6:** The table gives the list of 36 ED-AOPs that comprise the 7 connected components of the ED-AOP network along with the MIEs and AOs in each ED-AOP.

**Table S7:** The table lists the computed graph-theoretic measures including in-degree, out-degree, eccentricity and betweenness centrality, for the KEs in the two LCCs, C1 and C2. An event is said to be convergent if in-degree > out-degree for that particular event, while an event is divergent if in-degree < out-degree for that particular event, in the ED-AOP network.

**Table S8:** The list of 44 KEs in the LCC C1 categorized into 4 different systems-level perturbations namely, hepatic, metabolic, neurological, and reproductive, based on the associated information in AOP-Wiki on the perturbed cell types, organs or biological processes.

**Table S9:** The table provides the list of 35 chemical stressors associated with events in LCC C1. Following a comparison with DEDuCT 2.0, 16 of the 35 chemical stressors were identified as potential EDCs.

